# Enhanced efficiency in the bilingual brain through the inter-hemispheric cortico-cerebellar pathway in early second language acquisition

**DOI:** 10.1101/2023.11.16.567455

**Authors:** Zeus Gracia-Tabuenca, Elise B. Barbeau, Shanna Kousaie, Jen-Kai Chen, Xiaoqian Chai, Denise Klein

**Affiliations:** Department of Statistical Methods, University of Zaragoza, Zaragoza, Aragon, Spain; Department of Neurology and Neurosurgery, McGill University, Montreal, Quebec, Canada; School of Psychology, University of Ottawa, Ottawa, Ontario, Canada

**Keywords:** Bilingualism, Connectomics, Cerebellum, Functional Networks

## Abstract

The bilingual experience has a profound impact on the functional and structural organization of the brain, but it is not yet well known how this experience influences whole-brain functional network connectivity. We examined a well-characterized large sample (151 participants) of monolinguals and bilinguals with varied age of second language acquisition, who underwent resting-state functional magnetic brain imaging. We constructed comprehensive functional brain networks for each participant, encompassing cortical, subcortical, and cerebellar regions of interest. Whole-brain analyses revealed that bilingual individuals exhibit higher global efficiency than monolinguals, indicating enhanced functional integration in the brain. Moreover, the age at which the second language was acquired correlated with this increased efficiency, suggesting that earlier exposure to a second language has lasting positive effects on brain functional organization. Further investigation through the network-based statistics (NBS) approach indicates that this effect is primarily driven by heightened functional connectivity between association networks and the cerebellum. This work shows that early learning enhances global whole-brain efficiency and that the timing of learning of two languages has an impact on functional brain organization.

**Significance statement:** Long-term learning impacts brain organization at different spatial scales, and this may be particularly enhanced during early stages of life. Bilingualism offers a unique opportunity to test long-term learning effects in the human brain, given that exposure to a second language can occur from birth or later in life, and can be maintained over long periods of time. We found that second language acquisition in early childhood (before five years of age) enhances brain network efficiency, and that this effect goes beyond the language and cognitive control regions, in fact, the interhemispheric cortico-cerebellar circuit plays a key role. This work shows that the timing of bilingual learning experience alters the brain functional organization at the global and local levels.

## 1. Introduction

Bilingualism provides a window into questions about brain organization and language development. The brain demonstrates a remarkable capacity to undergo structural and functional change in response to experience throughout the lifespan. Evidence suggests that, in many domains of skill acquisition, the manifestation of this neuroplasticity depends on the age at which learning begins. Bilingualism provides an optimal model for discerning differences in how the brain wires when a skill is acquired from birth, when the brain circuitry for language is being constructed, and later in life, when the pathways subserving the first language are already well developed. Acquisition of a second language (L2) may have a positive impact on working memory (Kousaie et al, 2021; Kwon et al., 2021), cognitive control (Kousaie et al., 2017), facilitate perception of speech in noise (Kousiae et al, 2019), improve attention (Vega-Mendoza et al., 2015), may bolster cognition in aging (Bialystok et al., 2014), or even imply better prognosis after a brain injury (Alladi et al., 2015) or provide a potential protective benefit in epilepsy (Stasenko et al., 2023). Although, this impact on behavior is influenced by factors such as chronological age or type of task (Gunnerud et al., 2020; Ware et al., 2020), or learning environment (Gullifer et al., 2018), there is significant evidence that bilingualism shapes the brain at the functional (Berken et al., 2016a; Dash et al., 2022) and structural level (Luk et al., 2011; Klein et al, 2014; García-Pentón et al., 2014; Fedeli et al., 2021), and that these effects although most obvious when language is acquired in childhood, can even be observed when languages are learned later in life, and even over relatively short periods of time such as weeks (Barbeau et al. 2017). All these findings suggest that bilingualism, and second language learning, plays a role in the functional and structural organization of the brain.

Several studies have pointed out that functional changes related to bilingual processing are not restricted to language-related areas in the left frontal and temporal regions, but also to fronto-parietal systems involved in cognitive control (Li et al., 2015; Sulpizio et al., 2020; Kousaie et al 2017) and subcortical systems such as the basal ganglia involved in complex articulatory control (Klein et al, 1994; Klein et al, 1995; Klein et al 2006; Berken et al. 2015). There is also an emerging body of work showing that the cerebellum plays a relevant role in language (Booth et al., 2007; Murdoch, 2010; Fiez, 2016). Patients with lesions in the cerebellum have shown language-related deficits such as in grammar processing (Adamaszek et al., 2012) or in semantic fluency (Ahmadian et al., 2019). Functional neuroimaging studies during language tasks have found that activation (inferred by fMRI signal) mainly in the right posterolateral cerebellum is associated with performance on language tasks (Stoodley and Schmahmann, 2009; Stoodley, 2012; King et al., 2019), suggesting the relevance of the cortico-cerebellar system for optimal execution and processing. However, to date, it is not well understood how the cerebellar system impacts the acquisition of a second language at different ages of exposure, and in what way L2 acquisition influences the interactions of the cerebellar system with other cortical and subcortical brain regions.

Brain imaging research on bilingualism to date has generally focused on the role of specific brain areas (Berken et al., 2016a; Dash et al., 2022), on white matter tracts connecting language regions (Sander et al., 2022), and/or on specific resting-state functional networks such as the language and control networks (Gullifer et al. 2018). However, there is a substantial body of evidence showing that the brain is a complex network and that these network properties, modeled and inferred from graph analysis, can shed light on the general organizing principles of brain networks and how they relate to cognition (Rubinov and Sporns, 2010; Stam and Van Straaten, 2012). In particular, many studies have shown the modular organization of the brain networks, which reflects the segregation of different brain regions into specialized subsystems or modules (Meunier et al., 2010; Betzel et al., 2013; Puxeddu et al., 2020). Brain modularity has been proposed as a network-level biomarker of cognitive plasticity, the capacity to change and adapt cognitive performance (Gallen and D’Esposito, 2019). Concurrently, in addition to modularity or segregation, another critical feature of brain networks is their functional integration, which reflects how interconnected areas work together (Achard and Bullmore, 2007; Marrelec et al., 2008; Bullmore and Sporns, 2012). In particular, global efficiency, a graph coefficient (further described in the methods section) that accounts for the network’s integration, has been widely used to model functional connectomes and their relationship with cognition (Santarnecchi et al., 2014; Danti et al., 2018; Farah and Horowitz-Kraus, 2019), and with language performance (Pamplona et al., 2015). Higher global integration has been found to facilitate general cognitive ability (Wang et al., 2021).

Given the potential impact of second language learning on cognition and the wiring of the brain, it is of interest to study how bilingualism shapes the organization of the brain. To date, how integration and segregation of functional brain networks relate to early and late second language (L2) learning remains unknown. By examining a large sample of bilingual participants who differed in their ages of L2 acquisition, we were able to tease apart the effect of early versus late L2 learning on the wiring of the brain. We expect that early L2 learning enhances global whole brain efficiency and modularity. Brain network features were extracted from whole brain connectomes constructed from resting-state fMRI connectivity, encompassing cortical, subcortical, and cerebellar regions of interest. By investigating the whole-brain network and modularity and efficiency related to learning one versus two languages, we show how these effects are associated with learning one language or a second language early or later in life.

## 2. Methods

### 2.1 Sample

Resting-state fMRI data from 151 participants was compiled retrospectively from data collected in one laboratory (DK) using the same scanning parameters at the Montreal Neurological Institute (MNI). Subjects were grouped according to their exposure to English and French based on their self-reported age of acquisition (AoA) of their second language (L2). Groups included simultaneous bilinguals (AoA of both languages from birth), early bilinguals (L2 AoA before or equal to 5 years), late bilinguals (L2 AoA after 5 years), and monolinguals (limited exposure to an L2).

**Table 1.** Sample characteristics by mono/bilingual group in terms of the sample size, chronological age, sex distribution, age of L2 acquisition (AoA), years of education, and averaged in-scanner motion. Head-motion was estimated by the means of the subject-averaged framewise displacement (FD) in mm using the CONN’s standard algorithm. For numerical variables, mean +/- standard deviation is reported. Abbreviations: monolinguals (Mono), and late (Late.B), early (Early.B), and simultaneous (Sim.B) bilinguals.

|  | Mono | Late.B | Early.B | Sim.B |  |
| --- | --- | --- | --- | --- | --- |
| N | 19 | 59 | 34 | 39 |  |
| Age | 25.8±4.6 | 24.4±3.9 | 24.4±4.4 | 23.8±4.1 | $F_{(3,147)} = 1$ ($p = 0.39$) |
| Sex (F/M) | 12 / 7 | 34 / 25 | 27 / 7 | 24 / 15 | $\chi^2_{(3)} = 4.7$ ($p = 0.2$) |
| AoA | | 9.1±2.4 | 4.1±1.2 | 0 | $F_{(2,129)} = 342$ ($p < 2e-16$) |
| Education | 16.5±1.7 | 16±3.1 | 15.8±2.8 | 15.5±2.2 | $F_{(3,115)} = 0.6$ ($p = 0.63$) |
| Motion | 0.1±0.02 | 0.11±0.05 | 0.11±0.04 | 0.1±0.04 | $F_{(3,147)} = 0.4$ ($p = 0.74$) |

All participants self-reported good health, being right-handed, and null acquisition of any language besides English and French. Exclusion criteria included any language, hearing, or visual impairment, any history of traumatic brain injury, medical condition, or neurological disorder, and MRI incompatibility. All datasets came from studies approved by the Research Ethics Board at the MNI (McGill University) and all participants gave their written informed consent.

### 2.2 MRI session

Brain imaging was acquired on a 3-Tesla TrioTim Siemens scanner at the MNI using a 32-channel head coil. Resting-state fMRI consisted of T2-weighted echo-planar (EPI) sequenced 132 whole-brain volumes (TR/TE = 2260/30 ms; flip angle 90°; matrix size = 64 × 64; FoV = 224 mm; 38 3.5 mm thick slices) acquired in 5:04 minutes. Participants were instructed to keep their eyes open and to fixate on a cross that was presented at the center of the screen. For anatomical reference, high-resolution T1-weighted images were obtained from a 3D magnetization prepared rapid acquisition gradient echo (MPRAGE) sequence (TR/TE = 2300/2.98 ms; flip angle = 9°; matrix size = 256 × 256, FoV = 256 mm; slice thickness = 1 mm; interleaved excitation).

### 2.3 MRI preprocessing

Functional brain images were preprocessed following the standard pipeline of CONN toolbox v.20b (Whitfield-Gabrieli and Nieto-Castanon, 2012; Nieto-Castanon, 2020; RRID: SCR_009550) which relies on routines from SPM12 (Ashburner et al., 2014; RRID: SCR_007037). Briefly, fMRI data underwent realignment and unwarping, outlier identification, segmentation and normalization into MNI space, and smoothing with a 6 mm full width at half maximum (FWHM) Gaussian kernel. Outlier scans were identified using the conservative setting in ART (Artifact Detection Tools; RRID: SCR_005994, Nieto-Castanon, 2020; RRID: SCR_009550), which defines outliers as volumes in which average intensity deviated more than three standard deviations from the mean intensity or composite head movement exceeding 0.5 mm from the previous volume scan. In addition, potential confounding effects from physiological and other spurious sources of noise were estimated and regressed from the fMRI time series. Confounds included noise components from white matter and cerebrospinal fluid areas (aCompCor; Behzadi et al. 2007; Chai et al., 2012), estimated subject-motion parameters, and identified outlier scans (Satterthwaite et al. 2013).

### 2.5 Brain network features

For each participant, the preprocessed fMRI time series were averaged based on a brain segmentation of 374 regions of interest (ROI), which included 333 cortical areas (Gordon et al., 2016) plus 15 subcortical and 26 cerebellum areas (from CONN’s Harvard-Oxford template). These ROIs were encompassed in 14 modules: 12 predefined cortical modules, plus subcortical and cerebellum modules. Network edges were computed as the functional connectivity between every pair of ROIs, that is Pearson’s correlation of their corresponding pairwise ROI’s fMRI averaged signal. Additionally, the edge values underwent Fisher’s r-to-z transformation. Network organization features were assessed using the global efficiency (*E*) and modularity (*Q*) algorithms, which account for the functional integration and segregation of the networks, respectively (Rubinov and Sporns, 2010). Negative edges were discarded and weighted formulas were applied. *Q* was calculated using Newman’s algorithm (Clauset et al., 2004), which assesses the proportion of within-module connections across all possible connections. Higher values of *Q* express higher segregation of the network. *E* was computed as the average of all ROI’s nodal efficiency within each whole-network. Where nodal efficiency of a particular node (i.e., ROI in this case) accounts for the average of the inverted shortest paths to all other possible nodes. Shortest paths were computed following Dijkstra’s algorithm (Dijkstra, 1959). Then, higher values of *E* express higher network integration. Newman’s and Dijkstra’s algorithms were calculated via the R package ‘igraph’ (Csardi and Nepusz, 2006; RRID: SCR_019225). Formulas for *Q* and *E* are described in the Supplementary Information.

### 2.6 Sample inference statistics

Group effects were tested on the whole brain network using a one-way ANOVA, and Welch’s two-sample t-test (Welch, 1947) for each pair of groups as post hoc analysis. We corrected for multiple comparisons via the Bonferroni-Holm method (Holm, 1979). Age of L2 acquisition (AoA) effects were tested within the subset of early and late bilinguals by the means of Pearson’s correlation.

To understand the specific network connections contribution to the whole brain network segregation or modularity (Q and E) differences , we explored module level effects in those pairs of groups with significant *Q* and/or *E* whole-brain differences using a network-based statistics approach (NBS; Zalesky et al., 2010). NBS was used to identify clusters of connections below an a priori threshold and evaluate their cluster-level significance using a family-wise error (FWE) permutation test based on their sum of connectivity strengths. Furthermore, the NBS was applied recursively between the ROIs within the significant clusters of modules found. This approach was performed to determine which functional networks (modules) and/or particular brain regions are (or are not) driving the whole-brain effects. The NBS framework can effectively handle multiple comparison testing while preserving higher statistical power than traditional corrections, even with a greater number of regions of interest (ROIs) (Zalesky et al., 2010). NBS was calculated using the R library NBR v.0.1.5 (Gracia-Tabuenca and Alcauter, 2020; RRID: SCR_019114), setting the individual connectivity a priori threshold to p < 0.01 (two-sided) and running 1000 permutations. To test the consistency of the results, NBS analyses were performed additionally with an edgewise threshold of p < 0.05.

### 2.7 Additional covariates

Analyses were also implemented to check the potential effects of additional factors: years of experience (YoE) of the L2, and L2 proficiency, were related to the whole-brain network features (modularity and efficiency) and the functional connectivity between brain networks. YoE was calculated as the number of years the participant had been exposed to their L2, that is, chronological age minus AoA. For bilingual participants, L2 proficiency was rated using a self-report questionnaire with Likert-type responses ranging 1 to 7 regarding their current ability in their L2 on reading, speaking, writing, and comprehension, where 1 indicated “*very poor*” and 7 indicated “*native-like*” ability (Klein et al., 2014).

Given that cortico-cerebellar connectivity may be affected by other sensorimotor experiences, such as musical training (Shenker et al., 2023), we performed additional whole-brain and network analyses adding musical experience as a binary covariate (26 participants reported having musical training).

### 2.8. Code accessibility

The preprocessed data and the code/software described in the work is freely available online at https://github.com/zchuri/BilingualBrainNet

## 3. Results

### 3.1 Whole-brain efficiency was higher in early and simultaneous bilinguals compared to monolinguals

Significant group effects were found for the global efficiency feature (F_(3,147)_ = 3.39; p_FWE_ = 0.04). Post-hoc tests revealed that early and simultaneous bilingual groups, but not late bilinguals, had significantly higher global efficiency compared to the monolingual group (Figure 1; early: t_(40.3)_ = 3.26; p_FWE_ = 0.014; simultaneous: t_(42.7)_ = 2.77, p_FWE_ = 0.042; late t_(36.2)_ = 1.88, p_FWE_ = 0.272; Supplementary Table 1). Global efficiency did not differ significantly among the three bilingual groups (ps > 0.074). No group effects were found for the modularity feature (F_(3,147)_ = 0.53; p = 0.66). These effects were consistent using different connectivity density thresholds (Supplementary Figure 1). Furthermore, when testing the age of acquisition (AoA), we found a significant negative correlation with global efficiency (rho = -0.22; p-uncorrected = 0.037) (Supplementary Figure 2), but not with modularity (rho = -0.14; p = 0.18). This means that within the bilingual groups, the earlier the AoA the higher the network efficiency scores.

**Figure 1.**
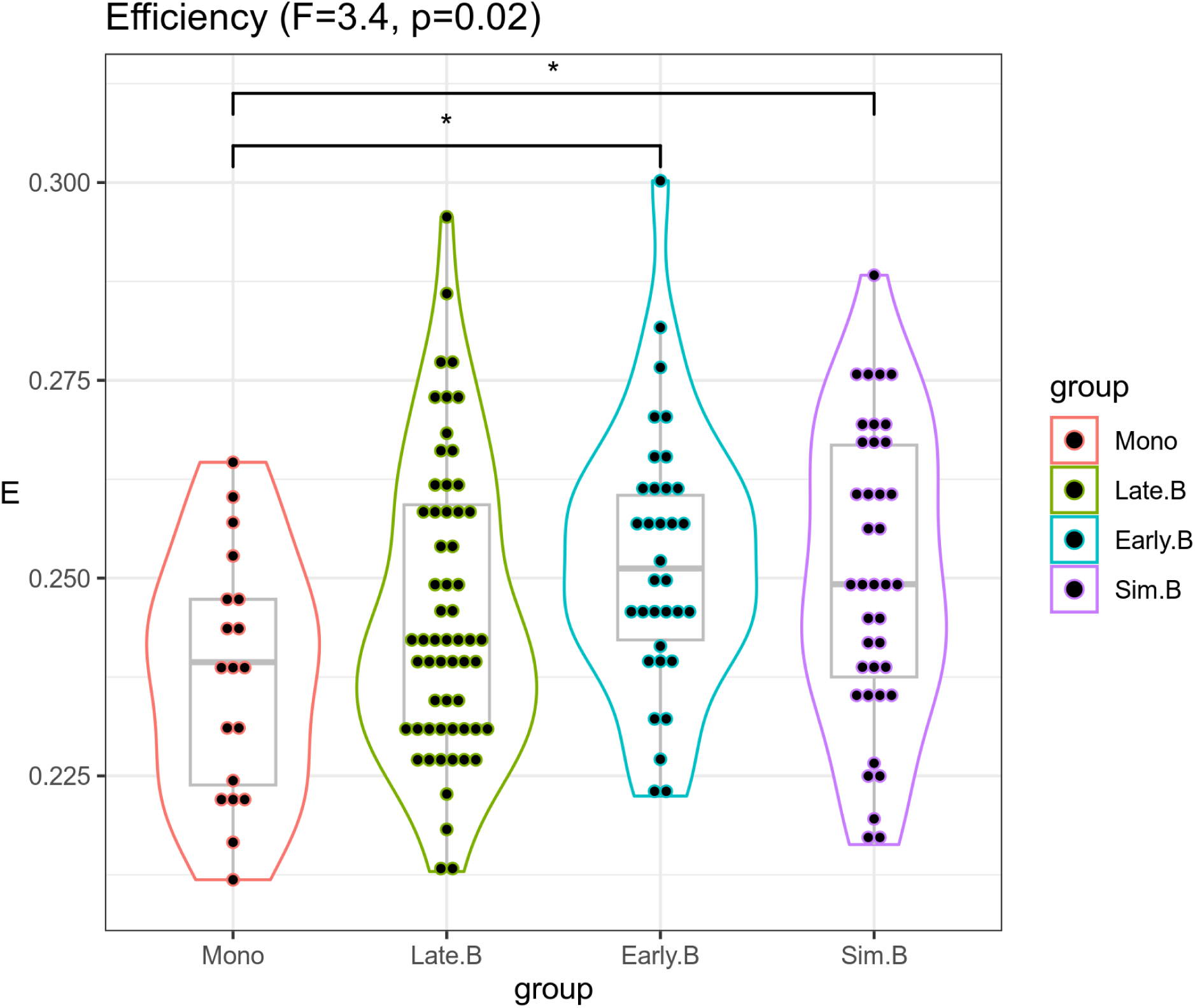
Group violin-boxplot for whole-brain global efficiency (E). ‘*’ stands for significance below p < 0.05 after Bonferroni-Holm correction. Group abbreviations: monolinguals (Mono), and late (Late.B), early (Early.B), and simultaneous (Sim.B) bilinguals.

### 3.2 Cortical and cortico-cerebellar connectivity was higher in early and simultaneous bilinguals compared to monolinguals: network-level results

We tested pairwise group differences between functional networks or modules using NBS between early/simultaneous bilinguals and monolinguals and found two significant (FWE-corrected) clusters of connections both with higher functional connectivity for the early and simultaneous bilingual groups compared to the monolinguals (Figure 2). Simultaneous bilinguals compared to the monolinguals exhibited higher connectivity between the cerebellum and a set of association networks including the default mode, dorsal and ventral attention, fronto-parietal, and sensorimotor-hand networks (p_FWE_ = 0.001; Figure 2; Supplementary table 2). In addition, connectivity between ventral attention and salience network was higher in simultaneous bilinguals compared to monolinguals. Early bilinguals compared to monolinguals again showed higher cerebellar connectivity with the association networks including default mode, dorsal attention, and fronto-parietal networks. Also, auditory network connectivity with the default mode and the ventral attention network was higher in early bilinguals compared to monolinguals (p_FWE_ = 0.001; Figure 2; Supplementary table 3).

**Figure 2.**
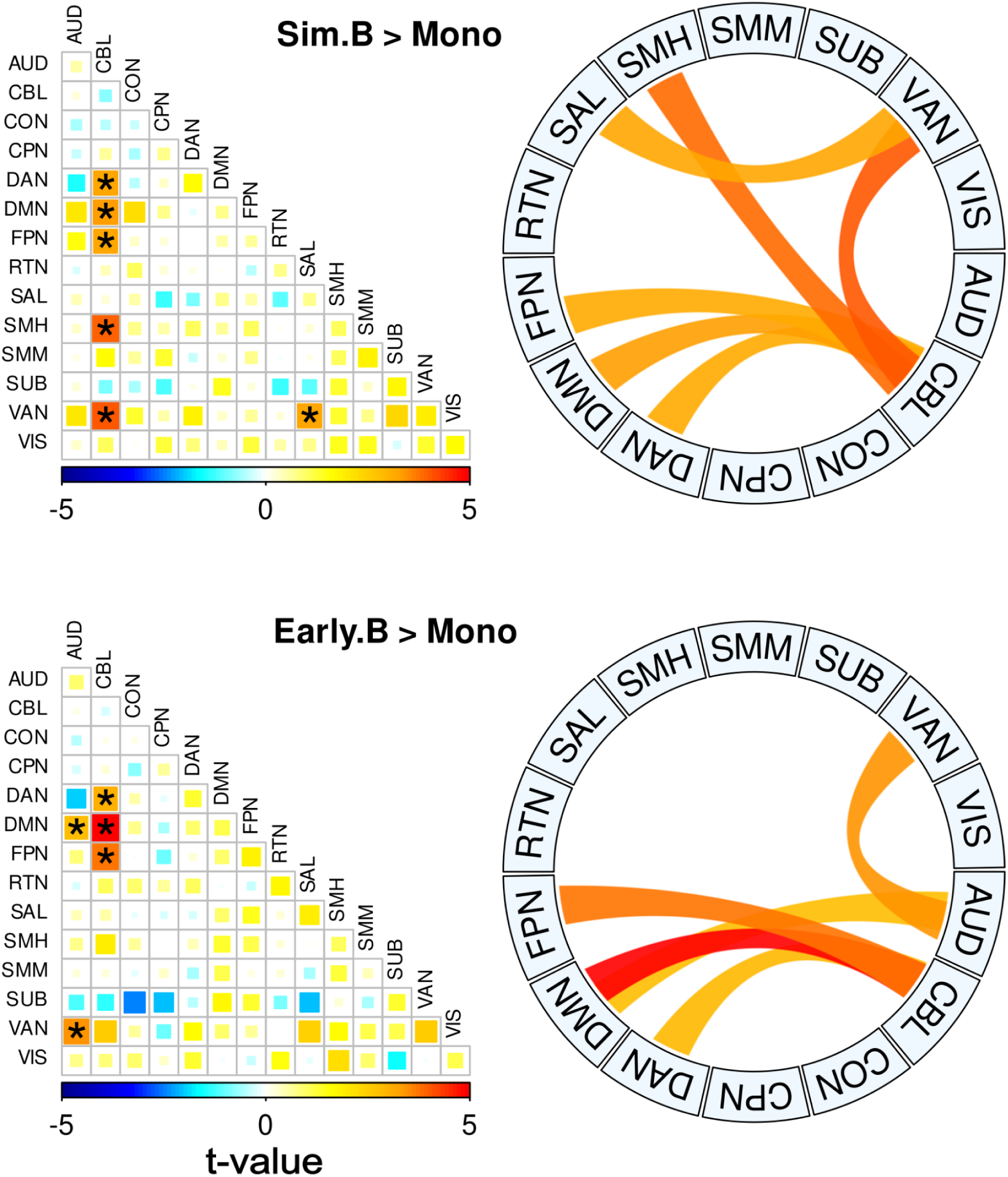
Pairwise plot and chord diagrams (Gu et al., 2014; RRID:SCR_002141) of significant (FWE-corrected) clusters of connections with higher functional connectivity between simultaneous (top) and early (bottom) bilinguals versus monolinguals. Abbreviations: AUD, auditory; CBL, cerebellar; CON, cingulo-opercular; CPN, cingulo-parietal; DMN, default mode; DAN, dorsal attention: FPN, fronto-parietal; RTN, retrosplenial-temporal; SAL, salience; SMH, sensorimotor-hand; SMM, sensorimotor-mouth; SUB, subcortical; VAN, ventral attention; VIS, visual.

### 3.3 Inter-hemispheric cortico-cerebellar connectivity was higher in simultaneous bilinguals: ROI-level results

We further tested the individual regions of interest within the previous network-wise clusters from section 3.2 and identified 118 connections that were stronger in simultaneous bilinguals compared to monolinguals (Figure 3). A significantly higher proportion of these connections were inter-hemispheric connections compared to intra-hemispheric connections (ratio = 64:37 = 1.72, excluding 17 connections involving vermis subregions; 𝞆2 = 7.22, p-value = 0.0072). In particular, the right cerebellum 1 lobules six and eight showed a higher number of inter-hemispheric connections (ten) to dorsal attention areas in the left hemisphere (Figure 3; Supplementary Table 4). NBS results were consistent with the additional p < 0.05 a priori threshold, same group contrasts at the module and ROI level resulted in FWE-corrected clusters, which encompassed the individual edges found at the p < 0.01 threshold (Supplementary Figure 3).

**Figure 3.**
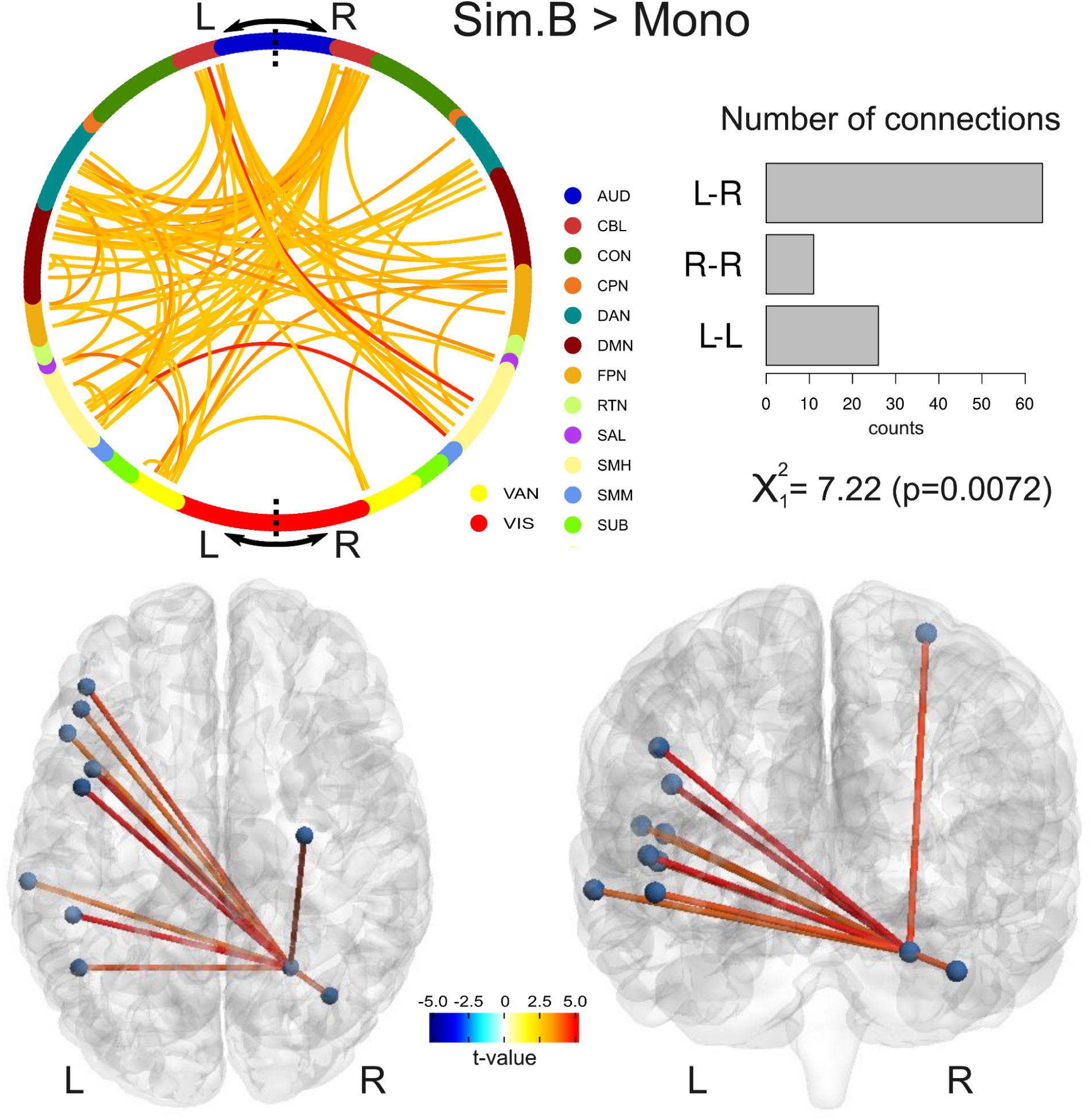
Regions-of-interest level results showing higher cortical and cortico-cerebellar connectivity in simultaneous than monolingual group. Chord diagram (Pedersen, 2020; RRID:SCR_021239) depicting a significant (FWE-corrected) cluster of edges with higher functional connectivity in the simultaneous compared to the monolingual group (top left). Nodes from each module are represented with different colors. Frequency barplot divided into intra-right (R-R), intra-left (L-L), or inter- (L-R) hemispheric edges with its corresponding proportion test (top right). Brain net (Xia et al., 2013; RRID:SCR_009446) representation of the corresponding right cerebellar lobule six edges dominated by interhemispheric connections (bottom).

### 3.4 Covariate effects

No significant relationships were found between L2 proficiency nor years of experience (YoE) and global efficiency or modularity, nor for functional connectivity between different networks from the NBS analysis. Regarding the music training covariate, significant group differences remain at the whole-brain and functional network levels when this factor is taken accounted for (Supplementary figure 4).

## 4. Discussion

Building on previous findings which suggest that bilingual language experience confers unique patterns of neurofunctional activity and brain structure (e.g., Mechelli et al., 2004; Klein et al, 2014; García-Pentón et al., 2014; Berken et al 2016a; Kousaie et al 2017; Fedeli et al., 2021; Dash et al., 2022), we explored the global impact of language learning by applying whole-brain network analyses on one of the largest well-characterized bilingual samples with neuroimaging data acquired on the same scanner with the same scanning parameters. We asked whether learning a second language either early or later in life, as compared to learning one language, affects the organization of whole-brain functional networks. A comparison of the influence of language learning experience on how the brain wires both in early childhood and later in life provides immeasurable insight into how learning experience influences brain structure and function and how the brain maximizes the efficiency of information processing (Watts and Strogatz, 1998; Achard and Bullmore, 2007; Butz et al., 2014). Simultaneous bilinguals constitute an apt point of comparison to monolinguals (similarly exposed to language from early life), hence, permitting the ideal basis for comparing language representation in the bilingual versus monolingual brain. Moreover, inclusion of individuals who ranged in age of L2 acquisition allowed us to distinguish the neurological correlates of simultaneous or early bilingualism as compared to sequential or late bilingualism.

Our results show that the global efficiency of functional connectomes in the bilingual groups is higher compared to the monolingual group, but when testing network modularity no significant effects are found. This suggests that L2 learning has little impact on the modular segregation of the brain subsystems, and that in fact how these modules interact is what matters with respect to brain organization. The findings are in agreement with an emerging body of evidence that shows that brain regions do not function in isolation, but rather how brain networks of different regions interact with one another is essential to achieve complex language comprehension, production, and supporting functions (Fedorenko et al., 2014; Hagoort, 2014; Chai et al., 2016; Sander et al., 2023). Moreover, brain regions and networks outside of the classic perisylvian areas are important for efficient language function and learning. More specifically, the higher global efficiency in simultaneous and early bilinguals was largely driven by cerebellar-cortical connections.

Our work extends previous brain imaging studies in bilingual participants by taking into consideration the interactions between cerebellum and the cortex, using a large bilingual sample, with participants who differ in age of second language acquisition. This varied L2 age of acquisition is critical to answer the question of how learning experience impacts how the brain wires, when the brain circuitry for language is being constructed, and later in life, when the pathways subserving the first language are already well developed. Previous work examining resting-state fMRI in bilingual samples has been particularly focused on how bilingual experience affects the functional connectivity of language and/or executive control networks (Pliatsikas and Luk, 2016). For instance, Berken et al. (2016a) found that simultaneous bilinguals have higher functional connectivity in bilateral inferior frontal gyrus (IFG), and higher connectivity between the IFG and the inferior parietal lobule (IPL) bilaterally, as well as posterior cerebellum, as compared to late bilinguals. Li et al. (2015) showed greater dorsal anterior cingulate functional connectivity with the left superior temporal sulcus in a group of sign-language bilinguals compared to a group of monolinguals. Gullifer et al. (2018) also found that early AoA correlates with higher interhemispheric frontal functional connectivity, and Li et al. (2015) showed lower functional connectivity within attention-related networks correlates with higher proficiency. To our knowledge, only one previous study attempted to address the bilingual brain from a whole-brain network perspective, Fan et al. (2021), did not find whole-network effects between a relatively small group of Mandarin monolinguals when compared to Cantonese-Mandarin bilinguals. However, it should be noted that these authors did not include the cerebellum in their analyses. Moreover, Cantonese and Mandarin share the same writing system and vocabularies but have more differentiation in phonology. Nevertheless, Fan et al (2021) did find that bilinguals had greater functional connectivity between the bilateral frontoparietal and temporal regions which is broadly consistent with our findings. We found higher whole-network efficiency in the bilingual brain, and differences in functional connectivity beyond the previously studied language and fronto-parietal networks, with the bilingual group showing higher connectivity between cerebellar and association cortical regions, in addition to higher connectivity between ventral attention and salience networks. Of note, our study demonstrates the major contribution of the cerebellar-cortical interaction in the organization of the bilingual brain. Importantly, we show that the age of acquisition of a second language (L2) plays a critical role on the functional organization of the brain – learning a second language during early childhood may result in more efficient wiring or more integration of brain networks, and stronger connections of the inter-hemispheric cortico-cerebellar pathway.

Studies utilizing tracing techniques have revealed that the cerebellar projections extend beyond the motor regions and into a greater extent of the cerebral cortex (Strick et al., 2009). While using resting-state fMRI to describe undirected pathways, Xue et al. (2021) were able to identify functional connections between the entire cerebellar cortex and both primary and association functional networks, including the vermis. Specifically, the contribution of the cerebellum to language learning has become increasingly apparent in the last few decades, as evidenced by a growing body of research (Mariën et al. 2014; Fiez, 2016). Studies describing patients with lesions in the cerebellum have reported impairments in prosody, grammar, syntax, and verbal fluency, among other functions (Schmahmann, 2019). Moreover, task-fMRI studies have consistently reported the simultaneous activation of the right posterior cerebellum and left frontal regions in language tasks (Stoodley, 2012; Keren-Happuch et al., 2014; Van Overwalle et al., 2014; Guell et al., 2018; King et al., 2019), with previous findings of highlighting right cerebellar-left frontal involvement in language learning, and particularly related to bilingual experience (Klein et al, 1995).

In addition, our results show that simultaneous bilinguals tend to have higher between-network functional connectivity and, particularly, in an inter-hemispheric fashion. This finding goes along with previous research showing higher interhemispheric effects in structural and functional neuroimaging features. For instance, Felton et al. (2017) found that bilinguals showed greater thickness in the corpus callosum compared to monolinguals, and DeLuca et al. (2019) found that fractional anisotropy in the corpus callosum correlated with AoA. On the other hand, early AoA has been associated with higher interhemispheric frontal functional connectivity (Berken et al., 2016a; Gullifer et al., 2018). What’s more, interhemispheric functional connectivity predicts prospective L2 acquisition performance (Sander et al., 2023). Although there is a lack of bilingual studies showing the intrinsic cortico-cerebellar connectivity, Krienen and Buckner (2009) used resting-state fMRI in an adult sample to identify four well-defined cortico-cerebellar circuits including motor and non-motor association regions, which showed greater contralateral functional connectivity.

## 5. Conclusion

We used network-based analyses in a large sample of monolinguals and bilinguals with different ages of L2 acquisition and identified whole-brain, network-level effects, as well as ROI-to-ROI interactions, including a high proportion of cortical and cerebellar-cortical interhemispheric connections related to L2 AoA. Bilingual brain networks showed greater efficiency that was driven by L2 exposure during early childhood. Taken together, these findings highlight how the brain’s intrinsic functional patterns are influenced by the developmental timeline in which second language acquisition occurs, and point to a more optimized mechanism to achieve L2 skillfulness, when language is learned during the period of significant brain growth and neuroplastic potential. As Berken et al (2016b) discuss, at the microscopic level, an enriched bilingual environment during the neonatal period may result in a cascade of biochemical events that increase production of the cellular substrates that regulate neuroplasticity, as well as the duration of their synthesis. This in turn, might result in macrostructural changes that manifest as efficient activation during speech, increased size of certain brain-language areas, and stronger connections between distributed brain regions within the language network. The degree to which these changes can occur depends on the age of second language exposure. When exposure occurs after the optimal periods for language acquisition are closing or have closed, neuroplasticity still occurs, allowing for L2 acquisition throughout the lifespan. However, the mechanisms for such neuro-plasticity later in life are likely to be qualitatively and quantitatively different from those biologically programmed to begin in early childhood. This work further enhances our understanding of how the timing of language experience affects brain efficiency and organization.

## Acknowledgments

Supported by grants from the Natural Sciences and Engineering Research Council of Canada (NSERC) (RGPIN-201405371) to D. Klein, the Blema and Arnold Steinberg Family Foundation, and funds from the CRBLM via the FRQNT/SC. ZGT was partially funded by the European Union’s NextGeneration programme and the Ministry of Universities RD 289/2021 UNI/551/2021 (Margarita Salas).

