## Supplementary Information for "Enhanced efficiency in the bilingual brain through the inter-hemispheric cortico-cerebellar pathway in early second language acquisition"

Formula 1: Newman's modularity is defined as (from package 'igraph' v.1.3.5 manual): "The modularity of a graph with respect to some division (or vertex types) measures how good the division is, or how separated are the different vertex types from each other. It defined as:"

$$Q = \frac{1}{2m} \sum_{i,j} \left( A_{ij} - \frac{k_i k_j}{2m} \right) \delta\left(c_i, c_j\right)$$

Where *m* is the number of edges, $A_{ij}$ is the element of the *A* adjacency matrix in row *i* and column *j*, $k_i$ and $k_j$ is the sum of weights of adjacent edges for nodes (vertex) *i* and *j*, respectively. $c_i$ and $c_j$ accounts for the module of *i* and *j*, respectively. And, $\delta\left(c_i, c_j\right)$ is 1 if $c_i = c_j$ and 0 otherwise.

Formula 2: Global efficiency is defined as the average nodal efficiency, that is, the network average of the inverse of the nodal shortest path distance:

$$E = \frac{1}{n} \sum_i e_i = \frac{1}{n} \sum_i \frac{\sum_{i \neq j} d_{ij}^{-1}}{n-1}$$

Where *n* accounts for the number of nodes in the network, $e_i$ is the efficiency of node *i*, and $d_{ij}$ is the distance between nodes *i* and *j* based on the weighted Dijktra's algorithm.

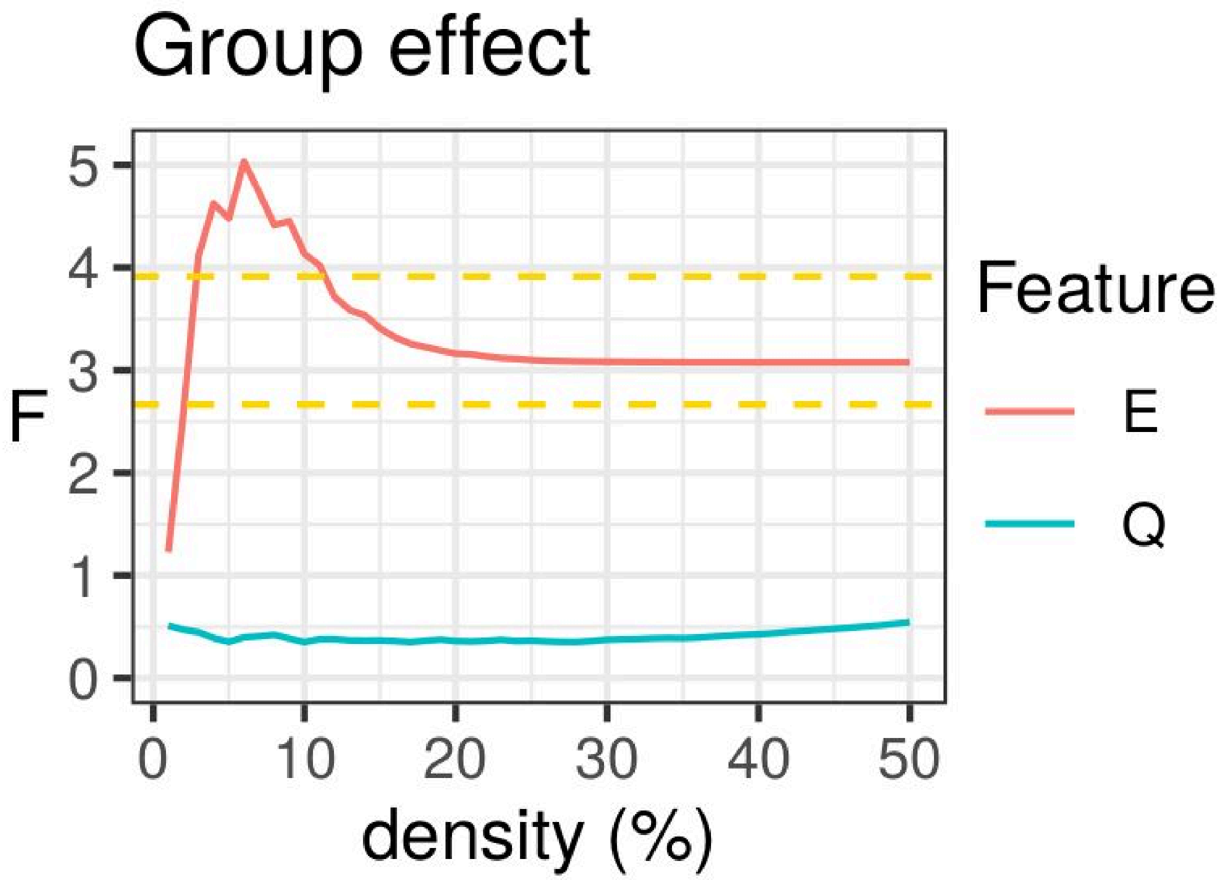


Supplementary Figure 1. Group effect of brain network features by connectivity density. Dashed yellow lines stand for significance of the F-test at p<0.05 and p<0.01. Feature abbreviations: efficiency (E) and modularity (Q).

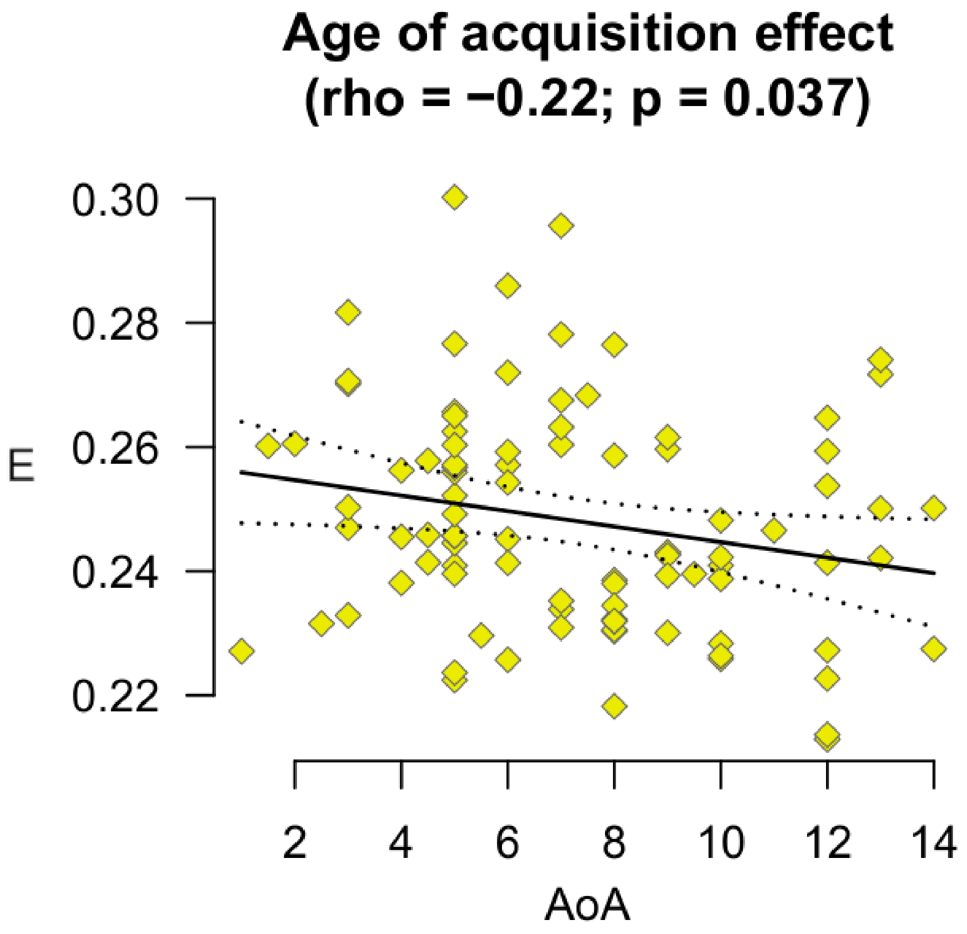


Supplementary Figure 2. Scatterplot with the relationship between whole-brain efficiency (E) and age of acquisition (AoA) in years for early and late bilingual participants.

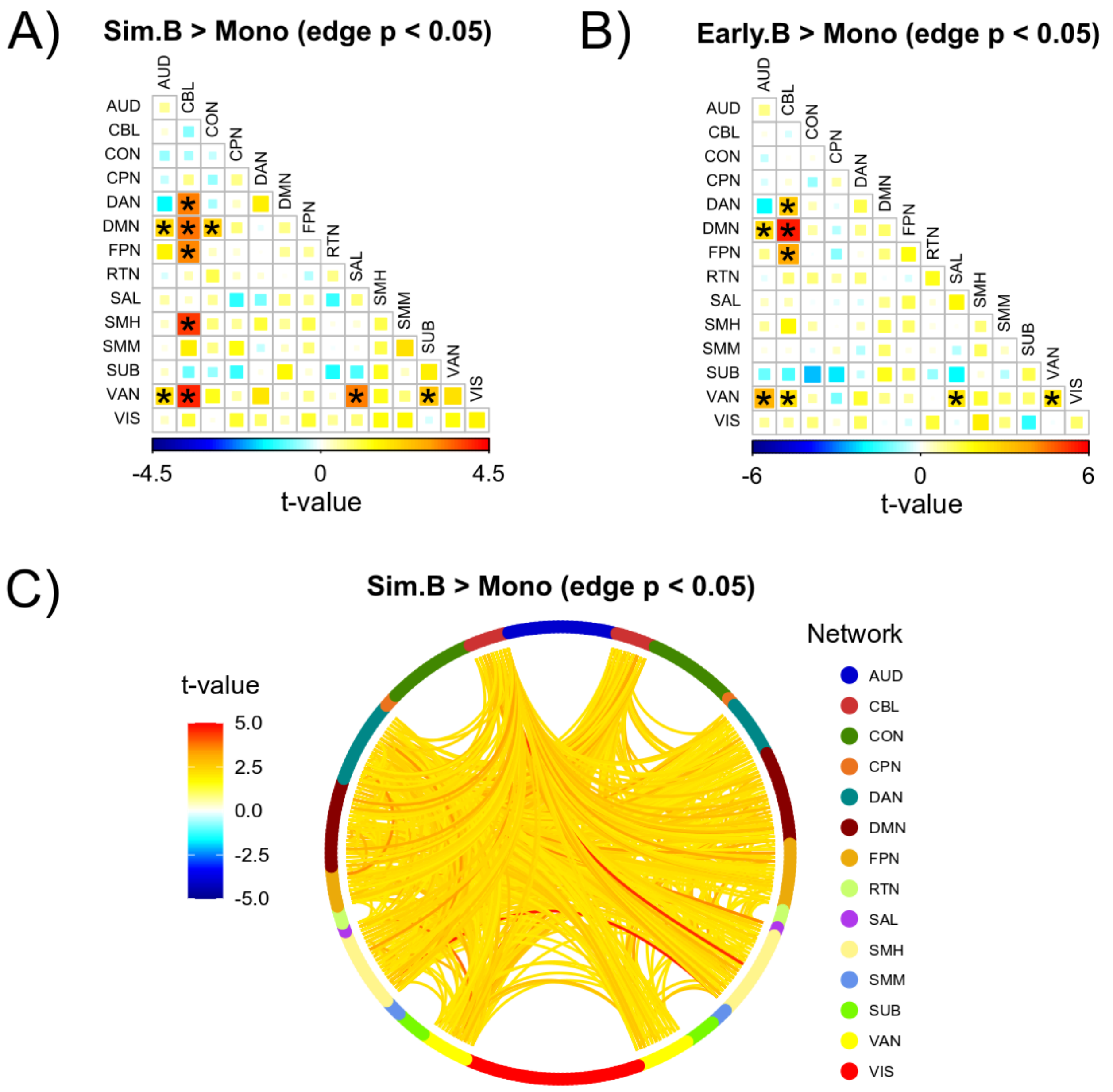


Supplementary Figure 3. NBS results with an edgewise a priori threshold of $p < 0.05$. Pairwise plots of significant (both $p = 0.001$; FWE-corrected) clusters of connections with higher functional connectivity between simultaneous (A) and early (B) bilinguals versus monolinguals at the module level. Chord diagram depicting a significant ($p = 0.001$; FWE-corrected) cluster of edges with higher functional connectivity in the simultaneous compared to the monolingual group at the ROI level (C). Abbreviations:

AUD, auditory; CBL, cerebellar; CON, cingulo-opercular; CPN, cingulo-parietal; DMN, default mode; DAN, dorsal attention: FPN, fronto-parietal; RTN, retrosplenial-temporal; SAL, salience; SMH, sensorimotor-hand; SMM, sensorimotor-mouth; SUB, subcortical; VAN, ventral attention; VIS, visual.

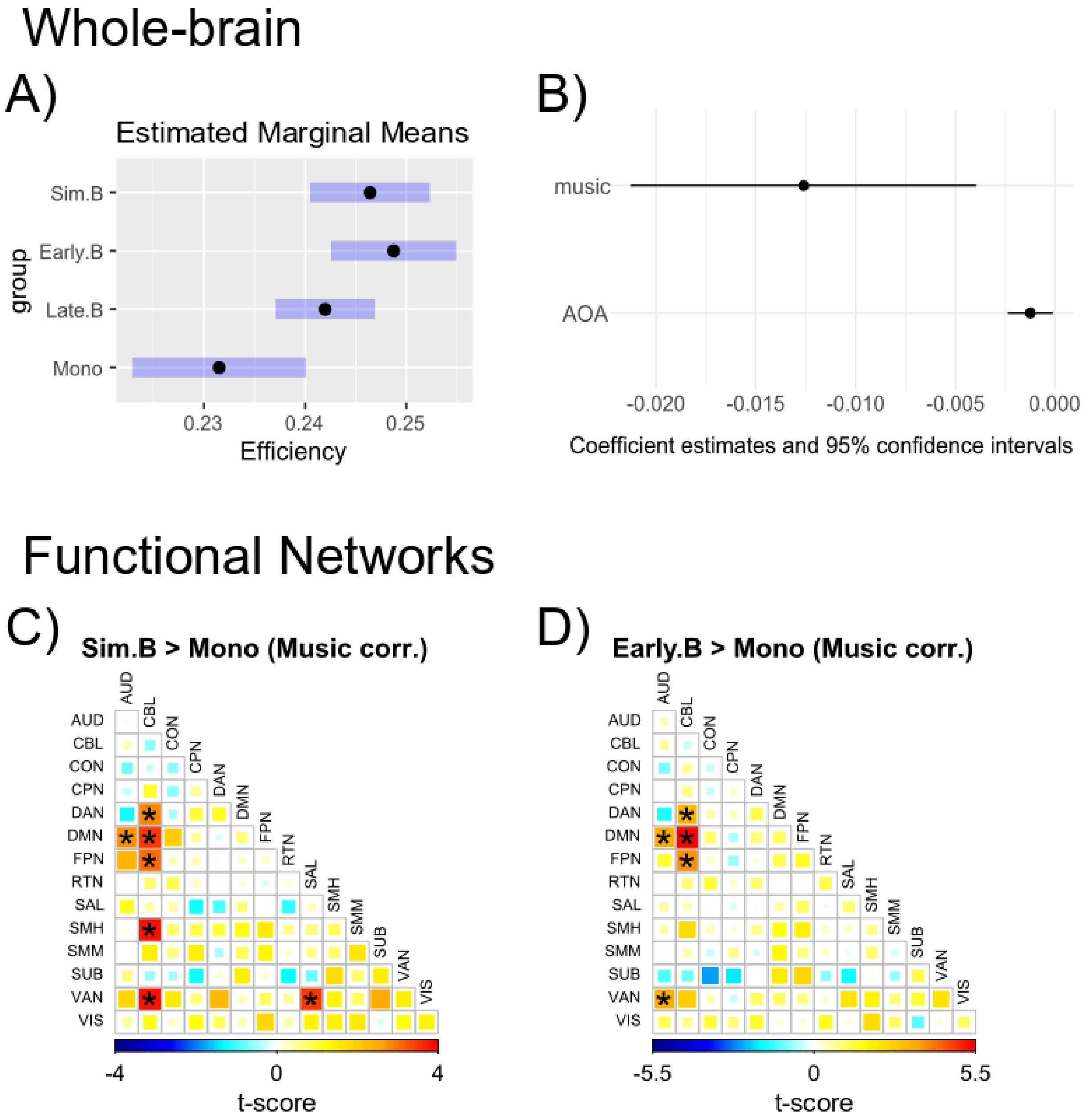


Supplementary Figure 4: Music training as binary covariate as reported in 26 participants. Top: Whole-brain global efficiency (*E*) Estimated Marginal Means (EMMs) for the mono/bilingual groups with music training as covariate (A). Estimated coefficients for the Age of Acquisition (AoA) and music covariate linear effects to E (B). Bottom: Pairwise plots of significant (FWE-corrected) clusters of connections with higher functional connectivity between simultaneous (C) and early (D) bilinguals versus monolinguals including binary music training as covariate. Abbreviations: AUD, auditory; CBL, cerebellar; CON, cingulo-opercular; CPN, cingulo-parietal; DMN, default mode; DAN, dorsal attention: FPN, fronto-parietal; RTN, retrosplenial-temporal; SAL, salience; SMH, sensorimotor-hand; SMM, sensorimotor-mouth; SUB, subcortical; VAN, ventral attention; VIS, visual. Corresponding 95% confidence intervals are depicted for A) and B) estimates.

Supplementary Tables

Supplementary table 1.

Post-hoc pairwise Welch's two-sample t-test between mono/bilingual groups for whole-brain global efficiency. Group abbreviations: monolinguals (Mono), and late (Late.B), early (Early.B), and simultaneous (Sim.B) bilinguals. Family-wise Error (FWE) correction was based on the Bonferroni-Holm method.

| **Contrast** | **t-score** | **DOF** | **p** | **pFWE** |
|---|---|---|---|---|
| Late.B - Mono | 1.88 | 36.2 | 0.068 | 0.222 |
| Early.B - Mono | 3.26 | 40.3 | 0.002 | 0.014 |
| Sim.B - Mono | 2.77 | 42.7 | 0.008 | 0.042 |
| Early.B - Late.B | 1.81 | 74.3 | 0.074 | 0.222 |
| Sim.B - Late.B | 1.24 | 80.8 | 0.218 | 0.436 |
| Sim.B - Early.B | -0.49 | 70.9 | 0.625 | 0.625 |

Supplementary table 2.

List of functional network (modules) connections with higher functional connectivity in the Simultaneous bilingual compared to the monolingual group within the FWE-corrected cluster using Network-Based Statistics. Network abbreviations: CBL, cerebellar; DMN, default mode; DAN, dorsal attention: FPN, fronto-parietal; SAL, salience; SMH, sensorimotor-hand; VAN, ventral attention.

| **Net 1** | **Net 2** | **t-value** | **DOF** | **p-value** |
|---|---|---|---|---|
| CBL | DAN | 3.25 | 56 | 0.002 |
| CBL | DMN | 3.38 | 56 | 0.0013 |
| CBL | FPN | 3.21 | 56 | 0.0022 |
| CBL | SMH | 3.93 | 56 | 0.0002 |
| CBL | VAN | 4.09 | 56 | 0.0001 |
| SAL | VAN | 3.29 | 56 | 0.0017 |

Supplementary table 3.

List of functional network (modules) connections with higher functional connectivity in the Early bilingual compared to the monolingual group within the FWE-corrected cluster using Network-Based Statistics. Network abbreviations: CBL, cerebellar; DMN, default mode; DAN, dorsal attention: FPN, fronto-parietal; SAL, salience; SMH, sensorimotor-hand; VAN, ventral attention.

| **Net 1** | **Net 2** | **t-value** | **DOF** | **p-value** |
|---|---|---|---|---|
| AUD | DMN | 2.88 | 51 | 0.0058 |
| AUD | VAN | 3.49 | 51 | 0.001 |
| CBL | DAN | 3.04 | 51 | 0.0037 |
| CBL | DMN | 5.58 | 51 | 9.28e-7 |
| CBL | FPN | 3.79 | 51 | 0.0004 |

Supplementary table 4.

List of ROI-to-ROI connections with higher functional connectivity in the Simultaneous bilingual compared to the monolingual group within the FWE-corrected cluster using Network-Based Statistics. Network abbreviations: AUD, auditory; CBL, cerebellar; CON, cingulo-opercular; CPN, cingulo-parietal; DMN, default mode; DAN, dorsal attention: FPN, fronto-parietal; RTN, retrosplenial-temporal; SAL, salience; SMH, sensorimotor-hand; SMM, sensorimotor-mouth; SUB, subcortical; VAN, ventral attention; VIS, visual.

| ROI1 | ROI2 | net1 | net2 | t | df | p |
|---|---|---|---|---|---|---|
| Area_8BM_L | Cerebellum Crus1 Left | DMN | CBL | 3.23 | 56 | 0.0021 |
| Area_8Ad_L | Cerebellum Crus1 Left | DMN | CBL | 2.84 | 56 | 0.0062 |
| Area_10d_L | Cerebellum Crus1 Left | DMN | CBL | 3.03 | 56 | 0.0037 |
| Area_2_R | Cerebellum Crus1 Left | SMH | CBL | 2.84 | 56 | 0.0063 |
| Area_posterior_47r_R | Cerebellum Crus1 Left | FPN | CBL | 3.11 | 56 | 0.0029 |
| Area_8Ad_R | Cerebellum Crus1 Left | DMN | CBL | 2.79 | 56 | 0.0072 |
| Area_8B_Lateral_R | Cerebellum Crus1 Left | DMN | CBL | 2.74 | 56 | 0.0083 |
| Area_PGi_L | Cerebellum Crus1 Right | DMN | CBL | 2.75 | 56 | 0.0081 |
| Area_8BM_L | Cerebellum Crus1 | FPN | CBL | 4.36 | 56 | 5.7e-05 |

| | | | | | | |
|---|---|---|---|---|---|---|
| | Right | | | | | |
| Area_8BM_L | Cerebellum Crus1 Right | DMN | CBL | 3.44 | 56 | 0.0011 |
| Area_8Ad_L | Cerebellum Crus1 Right | DMN | CBL | 3.07 | 56 | 0.0033 |
| Area_STSd_posterior_L | Cerebellum Crus1 Right | VAN | CBL | 2.72 | 56 | 0.0088 |
| Area_PGs_L | Cerebellum Crus1 Right | DMN | CBL | 2.8 | 56 | 0.007 |
| Area_8C_L | Cerebellum Crus1 Right | DAN | CBL | 3 | 56 | 0.004 |
| Area_IFSp_R | Cerebellum Crus2 Left | FPN | CBL | 3.11 | 56 | 0.0029 |
| Area_PGi_L | Cerebellum Crus2 Right | DMN | CBL | 2.71 | 56 | 0.0088 |
| Area_8BM_L | Cerebellum Crus2 Right | FPN | CBL | 3.37 | 56 | 0.0014 |
| Area_8C_L | Cerebellum Crus2 Right | DAN | CBL | 2.85 | 56 | 0.006 |
| Dorsal_area_6_R | Cerebellum 4 5 Left | SMH | CBL | 4.53 | 56 | 3.1e-05 |
| Area_8BM_L | Cerebellum 4 5 Right | DMN | CBL | 3.42 | 56 | 0.0012 |
| Dorsal_area_6_R | Cerebellum 4 5 Right | SMH | CBL | 3.3 | 56 | 0.0017 |
| Anterior_IntraParietal_Area_L | Cerebellum 6 Left | SMH | CBL | 2.67 | 56 | 0.0098 |
| Area_PH_L | Cerebellum 6 Left | DAN | CBL | 2.69 | 56 | 0.0095 |
| Dorsal_area_6_R | Cerebellum 6 Left | SMH | CBL | 3.4 | 56 | 0.0013 |
| Area_6mp_R | Cerebellum 6 Left | SMH | CBL | 2.89 | 56 | 0.0055 |
| Area_2_R | Cerebellum 6 Left | SMH | CBL | 2.69 | 56 | 0.0094 |

| | | | | | | |
|---|---|---|---|---|---|---|
| Area_STSd_posterior_L | Cerebellum 6 Right | VAN | CBL | 3.73 | 56 | 0.00045 |
| Area_IFSa_L | Cerebellum 6 Right | DAN | CBL | 3.29 | 56 | 0.0017 |
| Area_44_L | Cerebellum 6 Right | VAN | CBL | 2.72 | 56 | 0.0088 |
| Area_45_L | Cerebellum 6 Right | VAN | CBL | 2.76 | 56 | 0.0078 |
| Area_PH_L | Cerebellum 6 Right | DAN | CBL | 3.13 | 56 | 0.0028 |
| Rostral_Area_6_L | Cerebellum 6 Right | DAN | CBL | 3.7 | 56 | 0.00049 |
| Area_IFJa_L | Cerebellum 6 Right | DAN | CBL | 4.2 | 56 | 9.7e-05 |
| Area_TE1_Middle_L | Cerebellum 6 Right | DMN | CBL | 2.73 | 56 | 0.0084 |
| Dorsal_area_6_R | Cerebellum 6 Right | SMH | CBL | 3.18 | 56 | 0.0024 |
| Cerebellum Crus1 Right | Cerebellum 6 Right | CBL | CBL | 3.13 | 56 | 0.0028 |
| Area_46_L | Cerebellum 7b Left | DAN | CBL | 2.68 | 56 | 0.0095 |
| Area_2_R | Cerebellum 7b Left | SMH | CBL | 2.98 | 56 | 0.0042 |
| Area_IFSp_R | Cerebellum 7b Left | FPN | CBL | 2.68 | 56 | 0.0096 |
| Area_8B_Lateral_R | Cerebellum 7b Left | DMN | CBL | 3.23 | 56 | 0.0021 |
| Lateral_Area_7A_L | Cerebellum 7b Right | SMH | CBL | 2.67 | 56 | 0.0098 |
| Area_6_anterior_L | Cerebellum 7b Right | SMH | CBL | 2.77 | 56 | 0.0077 |
| Medial_IntraParietal_Area_L | Cerebellum 7b Right | DAN | CBL | 3.06 | 56 | 0.0034 |
| Area_IFJa_L | Cerebellum 7b Right | DAN | CBL | 2.86 | 56 | 0.006 |
| Dorsal_area_6_R | Cerebellum 7b Right | SMH | CBL | 2.81 | 56 | 0.0069 |
| Area_2_R | Cerebellum 7b Right | SMH | CBL | 2.69 | 56 | 0.0095 |
| Area_5m_L | Cerebellum 8 Left | SMH | CBL | 3.3 | 56 | 0.0017 |
| Area_7PC_R | Cerebellum 8 Left | SMH | CBL | 3.21 | 56 | 0.0022 |

| | | | | | | |
|---|---|---|---|---|---|---|
| Area_2_R | Cerebellum 8 Left | SMH | CBL | 3.11 | 56 | 0.0029 |
| Lateral_Area_7A_L | Cerebellum 8 Right | SMH | CBL | 2.85 | 56 | 0.006 |
| Area_5m_L | Cerebellum 8 Right | SMH | CBL | 2.79 | 56 | 0.0073 |
| Primary_Motor_Cortex_L | Cerebellum 8 Right | SMH | CBL | 2.7 | 56 | 0.0091 |
| Area_2_L | Cerebellum 8 Right | SMH | CBL | 2.95 | 56 | 0.0046 |
| Area_IFSa_L | Cerebellum 8 Right | DAN | CBL | 3.06 | 56 | 0.0034 |
| Medial_IntraParietal_Area_L | Cerebellum 8 Right | DAN | CBL | 3.61 | 56 | 0.00066 |
| Rostral_Area_6_L | Cerebellum 8 Right | DAN | CBL | 3.08 | 56 | 0.0032 |
| Area_posterior_9-46v_L | Cerebellum 8 Right | FPN | CBL | 3.28 | 56 | 0.0018 |
| Area_2_R | Cerebellum 8 Right | SMH | CBL | 2.72 | 56 | 0.0087 |
| Area_STSd_anterior_R | Cerebellum 8 Right | DMN | CBL | 2.9 | 56 | 0.0053 |
| Area_PGs_L | Cerebellum 9 Right | DMN | CBL | 2.74 | 56 | 0.0082 |
| Area_PH_L | Cerebellum 10 Right | DAN | CBL | 3.09 | 56 | 0.0031 |
| Cerebellum 7b Right | Cerebellum 10 Right | CBL | CBL | 2.71 | 56 | 0.0089 |
| Area_9_anterior_L | Vermis 1 2 | DMN | CBL | 3.05 | 56 | 0.0035 |
| Area_55b_L | Vermis 3 | VAN | CBL | 2.79 | 56 | 0.0072 |
| Area_STSd_anterior_R | Vermis 3 | DMN | CBL | 2.99 | 56 | 0.0041 |
| Area_23d_L | Vermis 4 5 | DMN | CBL | 2.7 | 56 | 0.0091 |
| Medial_IntraParietal_Area_L | Vermis 4 5 | DAN | CBL | 2.87 | 56 | 0.0057 |
| Area_PH_L | Vermis 4 5 | DAN | CBL | 3.11 | 56 | 0.0029 |

| | | | | | | |
|---|---|---|---|---|---|---|
| Dorsal_area_6_R | Vermis 4 5 | SMH | CBL | 3.58 | 56 | 0.00072 |
| Superior_Frontal_Language_Area_L | Vermis 6 | DMN | CBL | 2.72 | 56 | 0.0088 |
| Area_1_R | Vermis 6 | SMH | CBL | 2.83 | 56 | 0.0064 |
| Area_8BM_R | Vermis 6 | DMN | CBL | 2.95 | 56 | 0.0046 |
| Primary_Motor_Cortex_L | Vermis 7 | SMH | CBL | 2.81 | 56 | 0.0068 |
| Rostral_Area_6_L | Vermis 7 | DAN | CBL | 2.94 | 56 | 0.0047 |
| Area_3a_R | Vermis 7 | SMH | CBL | 2.8 | 56 | 0.007 |
| Area_posterior_9-46v_L | Vermis 8 | FPN | CBL | 2.68 | 56 | 0.0097 |
| Area_3a_R | Vermis 8 | SMH | CBL | 2.7 | 56 | 0.0091 |
| Area_2_R | Vermis 8 | SMH | CBL | 2.94 | 56 | 0.0048 |
| Area_TemporoParietoOccipital_Junction_1_R | Vermis 8 | VAN | CBL | 2.75 | 56 | 0.0079 |
| Area_5m_L | Area_6_anterior_L | SMH | SMH | 2.78 | 56 | 0.0074 |
| Area_2_L | Area_PFt_L | SMH | DAN | 3.43 | 56 | 0.0011 |
| Area_5m_L | Area_2_L | SMH | SMH | 3.07 | 56 | 0.0033 |
| Area_Lateral_IntraParietal_ventral_L | Area_2_L | DAN | SMH | 2.71 | 56 | 0.0089 |
| Area_8BM_L | Area_posterior_47r_L | FPN | VAN | 2.85 | 56 | 0.006 |
| Superior_Frontal_Language_Area_L | Area_47s_L | VAN | VAN | 2.84 | 56 | 0.0063 |
| Area_44_L | Area_47s_L | VAN | VAN | 2.74 | 56 | 0.0082 |
| Area_8BM_L | Area_47l_(47_lateral)_L | FPN | VAN | 2.84 | 56 | 0.0062 |

| | | | | | | |
|---|---|---|---|---|---|---|
| Anterior_Agranular_Insula_Complex_L | Area_47l_(47_lateral)_L | SAL | VAN | 3.92 | 56 | 0.00025 |
| Area_2_L | Medial_IntraParietal_Area_L | SMH | DAN | 2.71 | 56 | 0.0089 |
| Area_2_L | Area_PH_L | SMH | DAN | 2.72 | 56 | 0.0086 |
| Area_2_L | Area_PH_L | SMH | DAN | 2.97 | 56 | 0.0043 |
| Area_TE1_posterior_L | Area_8C_L | FPN | DAN | 3.49 | 56 | 0.00096 |
| Area_45_L | Area_8C_L | VAN | DAN | 2.77 | 56 | 0.0076 |
| Anterior_Agranular_Insula_Complex_L | Area_IFJa_L | SAL | DAN | 2.97 | 56 | 0.0044 |
| Area_PH_L | Area_IFJa_L | DAN | DAN | 2.78 | 56 | 0.0073 |
| Area_posterior_9-46v_L | Area_IFJa_L | FPN | DAN | 2.67 | 56 | 0.01 |
| Area_8BM_L | Area_TE1_Middle_L | FPN | DMN | 2.91 | 56 | 0.0052 |
| Area_TE1_Middle_L | Area_9_anterior_L | DMN | DMN | 2.82 | 56 | 0.0066 |
| Area_8BM_L | Superior_6-8_Transitional_Area_R | FPN | DMN | 2.73 | 56 | 0.0084 |
| Area_8C_L | Area_TE1_posterior_R | DAN | FPN | 3.31 | 56 | 0.0016 |
| Area_47l_(47_lateral)_L | Area_8BM_R | VAN | FPN | 3.54 | 56 | 0.00082 |
| Superior_Frontal_Language_Area_L | Area_anterior_32_prime_R | DMN | SAL | 3.54 | 56 | 0.00082 |
| Area_9_Posterior_L | Area_anterior_32_prime_R | DMN | SAL | 2.81 | 56 | 0.0069 |
| Area_PH_L | Area_6_anterior_R | DAN | DAN | 3.29 | 56 | 0.0017 |
| Area_5m_L | Area_6mp_R | SMH | SMH | 2.87 | 56 | 0.0057 |

| | | | | | | |
|---|---|---|---|---|---|---|
| Area_7PC_R | Area_posterior_47r_R | SMH | FPN | 3.12 | 56 | 0.0029 |
| Superior_Frontal_Language_Area_L | Anterior_Agranular_Insula_Complex_R | DMN | VAN | 2.67 | 56 | 0.0099 |
| Area_47s_L | Anterior_Agranular_Insula_Complex_R | VAN | VAN | 2.72 | 56 | 0.0088 |
| Area_47l_(47_lateral)_L | Anterior_Ventral_Insular_Area_R | VAN | SAL | 2.86 | 56 | 0.006 |
| Area_5m_L | Medial_IntraParietal_Area_R | SMH | DAN | 2.81 | 56 | 0.0068 |
| Medial_IntraParietal_Area_L | Area_IntraParietal_0_R | DAN | DAN | 2.91 | 56 | 0.0051 |
| Area_PH_L | Area_IntraParietal_0_R | DAN | DAN | 2.92 | 56 | 0.005 |
| Area_2_R | Area_IntraParietal_0_R | SMH | DAN | 3.08 | 56 | 0.0032 |
| Area_PH_L | Area_PHT_R | DAN | DAN | 2.84 | 56 | 0.0062 |
| Area_2_L | Area_PFt_R | SMH | SMH | 4.65 | 56 | 2.1e-05 |
| Area_2_L | Rostral_Area_6_R | SMH | DAN | 2.81 | 56 | 0.0068 |
| Dorsal_area_6_L | Area_8BM_R | SMH | DMN | 2.91 | 56 | 0.0052 |
| Area_STSd_anterior_R | Auditory_5_Complex_R | DMN | VAN | 2.98 | 56 | 0.0042 |
